# Human ACE2 peptide mimics block SARS-CoV-2 Pulmonary Cells Infection

**DOI:** 10.1101/2020.08.24.264077

**Authors:** Philippe Karoyan, Vincent Vieillard, Estelle Odile, Alexis Denis, Amélie Guihot, Charles-Edouard Luyt, Luis Gómez-Morales, Pascal Grondin, Olivier Lequin

## Abstract

In the light of the recent accumulated knowledge on SARS-CoV-2 and its mode of human cells invasion, the binding of viral spike glycoprotein to human Angiotensin Converting Enzyme 2 (hACE2) receptor plays a central role in cell entry. We designed a series of peptides mimicking the *N*-terminal helix of hACE2 protein which contains most of the contacting residues at the binding site and have a high helical folding propensity in aqueous solution. Our best peptide mimics bind to the virus spike protein with high affinity and are able to block SARS-CoV-2 human pulmonary cell infection with an inhibitory concentration (IC_50_) in the nanomolar range. These first in class blocking peptide mimics represent powerful tools that might be used in prophylactic and therapeutic approaches to fight the coronavirus disease 2019 (COVID-19).

## Introduction

The coronavirus disease 2019 (COVID-19), caused by the severe acute respiratory syndrome-coronavirus 2 (SARS-CoV-2) has emerged as a pandemic, claiming at the time of writing more than 600,000 deaths and over 14.5 million confirmed cases world-wide between December 2019 and July 2020.^1^ Since SARS-CoV-2 discovery^2,3^ and identification, the energy deployed by the scientific community has made it possible to generate an extraordinary wealth of information. Whatever, to date, clinically approved vaccines or drugs are lacking.^4^ Indeed, no specific drugs targeting the virus are available^5^ and many clinical trials have been engaged with SARS-CoV-2 non-specific treatments.^6^ The structural and biochemical basis of infection mechanism has been investigated, highlighting that the virus cell-surface spike protein of SARS-CoV-2 is targeting human receptors.^4,7^ Human Angiotensin Converting Enzyme 2 (hACE2) and the cellular Transmembrane Protease Serine 2 (TMPRSS2) have been identified as major actors of the virus entry into human cells.

With the goal of preventing the SARS-CoV-2 from infecting human cells, blocking the interaction between hACE2 and the virus spike protein has been validated. Indeed, inhibition of SARS-CoV-2 infections in engineered human tissues using clinical-grade soluble ACE2 was recently demonstrated.^8^ Likewise, an engineered stable mini-protein mimicking three helices of hACE2 to plug SARS-CoV-2 spikes^9^ was described, but its capacity to block viral infection was not demonstrated. If several in silico designed peptides were proposed to prevent formation of the fusion core,^10^ first attempts to design a peptide binder derived from hACE2 proved to be a difficult task, leading to mitigated results.^11^

Thus, starting from the published crystal structure of SARS-CoV-2 spike receptor-binding domain (RBD) bound to hACE2,^12^ we designed peptide mimics of the *N*-terminal hACE2 helix which interact with spike protein. We report here the strategy implemented to optimize the design of our peptide mimic, its high helical folding propensity in water and its ability to block cells infection by SARS-CoV-2 with an IC_50_ in the nM range upon binding to spike RBD with strong affinity. We also demonstrate the non-toxicity of our mimic at concentrations 150 times higher than IC_50_ on pulmonary cell lines.

## Results

### Design of peptides mimicking helix H1 of hACE2

We first examined the complex between hACE2 and the surface spike protein of SARS-CoV-2 (PDB 6m0j)^12^ in order to highlight the important contacts and some relevant characteristics of the interacting hACE2 sequence (**Figure 1**).

**Figure 1.**
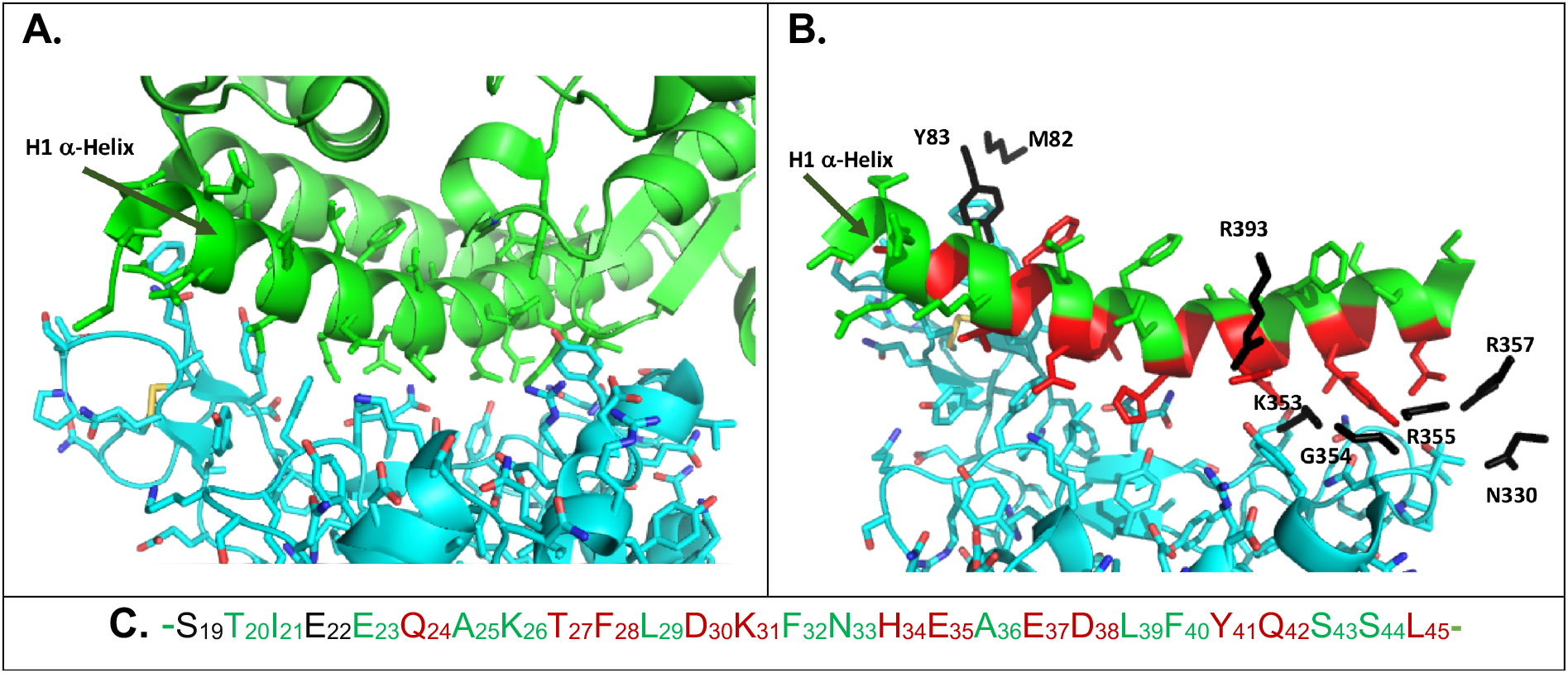
Structure of the complex between hACE2 and the spike protein of SARS-CoV-2 (pdb 6m0j). **1A**. hACE2 protein is shown in green and SARS-CoV-2 spike protein in cyan. **1B**. Residues of hACE2 interacting with spike protein are colored in red. Apart from **H1** helix, 8 additional residues from hACE2 are indicated in black. **1C. H1** helix sequence showing 12 interacting residues in red. Residue positions in green were considered as possible substitution sites for helical peptide design.

Twenty residues from hACE2 were identified^12^ as close contacts, using a distance cut-off of 4 Å. These interactions occur mainly through the *N*-terminal α-helix **H1** of hACE2 (**Figure 1A**). This α-helix, composed of 27 residues (from S19 to L45, **Figure 1C**) contains 12 residues (highlighted in red in **Figure 1B**) involved in hydrophobic interactions, hydrogen bonds and salt bridges.^12^

Our strategy was to design a peptide with high helical folding propensity and retaining most of the binding affinity of hACE2 to spike RBD of SARS-CoV-2, using natural amino acids^13^. Indeed, we preferred not using complex chemical tools known to stabilize α-helix^14,15^ in order to limit developability constraints. To design our mimics, we set up an optimization process with the dual aim of optimizing the helical content and limiting the antigenicity, to avoid triggering a neutralizing immune response that would compromise the peptide therapeutic potential. A combination of the Agadir program,^16^ an algorithm developed to predict the helical content of peptides, and the semi-empirical method reported by Kolaskar^17,18^ to highlight the number of antigenic determinants (AD), was iteratively used.

We observed that the *N*-terminal sequence of **H1** helix, composed of 4 residues (S_19_TIE_22_), corresponds to a consensus *N*-capping box motif (SXXE).^19^ A capping box features reciprocal backbone-side-chain hydrogen-bonds favoring helix initiation. Although this sequence does not adopt the H-bonded capping conformation in the crystal structure of the full protein, it could still be an important stabilizing element in the isolated helix when extracted from the protein context. These observations led us to keep 14 residues from the native **H1** helix of hACE2 as contact residues or putative stabilizing capping box. The 13 left residues that are not essential for the interaction were considered as possible sites for amino acid substitutions (**Figure 1C**). We thus substituted non-essential positions by Ala and/or Leu residues displaying higher helical folding propensities and calculated the peptide helical content after each substitution (**Table S1**).

A peptide sequence optimization was then realized to lower the antigenicity while keeping the helical propensity thanks to an iterative residue scanning and calculation of the helical content variation upon new substitutions (**Table S2**). This strategy highlighted the influence of the residue N33 in the native sequence. Indeed, if the N33/L33 substitution systematically improved the helical content, it was always at the expense of antigenicity. Conversely, the L33/N33 substitution reduced the antigenicity at the expense of helical content. The solution was found by L33/M33 substitution decreasing the number of AD.

The **H1** helix of ACE2 adopts a kinked conformation in the crystal structure, leading to a distorted CO/HN hydrogen bond network between H34/D38 and E35/L39 residues. Therefore, we considered introducing a proline as this residue is known to induce local kinks or distortions in natural helices.^20,21^ D38 was classed as a contact residue, while L39 side chain is not involved in any direct interaction. Consequently, L39 position was selected for substitution by proline (peptide **P5**, **Table 1**).

**Table 1.** Sequences and properties of synthesized peptides.

| Code | Sequence<sup>a</sup> | Predicted helical content %<sup>b</sup> | Predicted antigenicity (AD)<sup>c</sup> | Experimental helical content %<sup>d</sup> | %inhibition of SARSCoV-2 replication at 10 μM<sup>e</sup> | IC<sub>50</sub> (μM)<sup>g</sup><br>VeroE6/Calu3 | Kd<sup>h</sup> nM |
| --- | --- | --- | --- | --- | --- | --- | --- |
| **P1** | STIEE QAKTF LDKFN HEAED LFYQS SL-NH<sub>2</sub> | 6 | 0 | 7 | 9 |  |  |
| **P1scr** | AHLFS YLTTK EEQDN DAIFL QEFKS ES-NH<sub>2</sub> | 1 | 0 | 8 | 7 |  |  |
| **Ppen** | IEE QAKTF LDKFN HEAED LFYQS-NH<sub>2</sub> | 1 | 0 | 10 | 14 |  |  |
| **P2** | SALEE QLKTF LDKFL HELED LLYQL SL-NH<sub>2</sub> | 64 | 1 | 70 | 16 |  |  |
| **P3** | SALEE QLKTF LDKFL HELED LLYQL AL-NH<sub>2</sub> | 78 | 1 | 64 | 4 |  |  |
| **P4** | Ac-SALEE QLKTF LDKFL HELED LLYQL AL-NH<sub>2</sub> | 66 | 1 | 70 | 24 |  |  |
| **P5** | SALEE QLKTF LDKFL HELED PLYQL AL-NH<sub>2</sub> | 34 | 1 | 17 | 27 |  |  |
| **P6** | SALEE QLKTF LDKFL HELED LLYQL ALAL-NH<sub>2</sub> | 88 | 1 | 80 | 6,5 |  |  |
| **P7** | SALEE QYKTF LDKFL HELED LLYQL ALAL-NH<sub>2</sub> | n.a. | n.a. | 63 | 54 | 7 / 6,8 |  |
| **P8** | SALEE QLKTF LDKFM HELED LLYQL AL-NH<sub>2</sub> | 68 | 0 | 70 | 88 | 0.8 / 0,09 | 15.9 ± 2 nM |
| **P9** | SALEE QYKTF LDKFM HELED LLYQL SL-NH<sub>2</sub> | n.a. | n.a. | 53 | 100<sup>f</sup> | 0.3 / 0,07 | <1 nM |
| **P10** | SALEE QYKTF LDKFM HELED LLYQL AL-NH<sub>2</sub> | n.a. | n.a. | 56 | 100<sup>f</sup> | 0.06 / 0,08 | <1 nM |

<sup>a</sup> Amino acid sequence colored as follows: red, contact residue; green, non-essential residue; blue, substituted residue with respect to native sequence. Homotyrosine residue is depicted with underlined letter **Y**.
<sup>b</sup> Predicted helical content using Agadir program; n.a., not applicable (for hTyr containing sequences).
<sup>c</sup> Predicted antigenicity calculated from the predicting antigenic peptide tool from the immunomedicine group<sup>23</sup>; n.a., not applicable (for hTyr containing sequences).
<sup>d</sup> Experimental helical content determined by circular dichroism (see **Figure 2**).
<sup>e</sup> % of inhibition on Vero-E6 cells of at least three independent experiments (see **Figure 3A**).
<sup>f</sup> For peptides **P9** and **P10**, inhibition of infection was measured in the range 0.1-1 μM.
<sup>g</sup> Determined on VeroE6 and Calu3 cells only for peptides able to inhibit SARSCoV-2/VeroE6 cell infection.
<sup>h</sup> Dissociation constants with SARS-Cov-2 RBD-SD1 measured by BLI for peptides with IC<sub>50</sub> in the nM range.

In order to increase the helical content to a maximum level, this iterative study was also applied to longer peptide sequences starting from the 29-mers native one, albeit at the expense of antigenicity. Diverse combinations of *N*- and *C*-terminus capping groups were also considered (free extremities or *N*-acetyl, *C*-carboxamide groups).

Finally, we examined the possibility of promoting additional side chain contacts provided by ACE2 residues that do not belong to **H1** helix. Y83 residue appeared as a good candidate as it lies very close in space to A25 in **H1** helix. Molecular modeling was carried out on **H1** helix analogs in which A25 was replaced by tyrosine or homotyrosine (hTyr) residues (**Figure S1**). Calculations showed that *h*Tyr residue was able to project the phenol ring in the adequate 3D space to mimic Y83 position. Of note, *h*Tyr is a natural amino acid.^22^

Three peptides were selected as controls in our optimization process, **P1** (native sequence), **P1scr** (Scrambled peptide from **P1**) and **Ppen** (described by Pentelute & al. in a longer biotinylated and pegylated construct and termed SBP1 as a putative spike binder).^11^

The results highlighting the progression in the helical content and the number of antigenic determinants is reported in **Table 1** for the most relevant peptide mimics (see table S1, S2 and S3 for all peptide mimics designed and/or synthesized). These peptides were synthesized on a 5 to 20 mg scale from Fmoc-protected amino acids utilizing standard solid phase peptide synthesis (SPPS) methods on rink amide resin (See Methods).

### The designed peptides highlight an excellent correlation between calculated and experimentally determined helical content by circular dichroism in aqueous media

The conformation of synthesized peptides in aqueous solution was investigated by CD spectroscopy.^24^ **Figure 2** shows the superimposed CD spectra of 12 peptides, including the control ones, *i.e.* **P1** (native sequence), **P1scr** (Scramble), **Ppen.** The CD spectra of peptides **P1** (native), **P1scr** (scrambled) are characteristic of a predominant random coil structure with a negative minimum near 200 nm, as expected. Similarly, **Ppen** described as a helical peptide sequence^11^ adopted also a random coil conformation in solution. For all other peptides, the CD spectra exhibit the canonical α-helix signature, with a double minimum around 208 and 222 nm, with the exception of the proline containing **P5** peptide. The deconvolution of the CD spectra using DICHROWEB allowed estimating the helical population for each peptide, which is reported in **Table 1**. Overall, an excellent agreement was observed between the AGADIR-computed values and the experimental helical population inferred from CD data. The native hACE2 H1 helix sequence (peptides **P1**, **Ppen**) has a weak propensity to fold into an α-helix in aqueous solution (below 10%). In contrast, sequence optimization lead to **H1** analogs exhibiting a high helical propensity (between 50 and 80%). The introduction of a proline residue in peptide **P5** has a strong destabilizing effect on helical conformation (17%) whereas L/hY substitution only led to a slight decrease of helical content (**P7** versus **P6** and **P10** versus **P8**).

**Figure 2.**
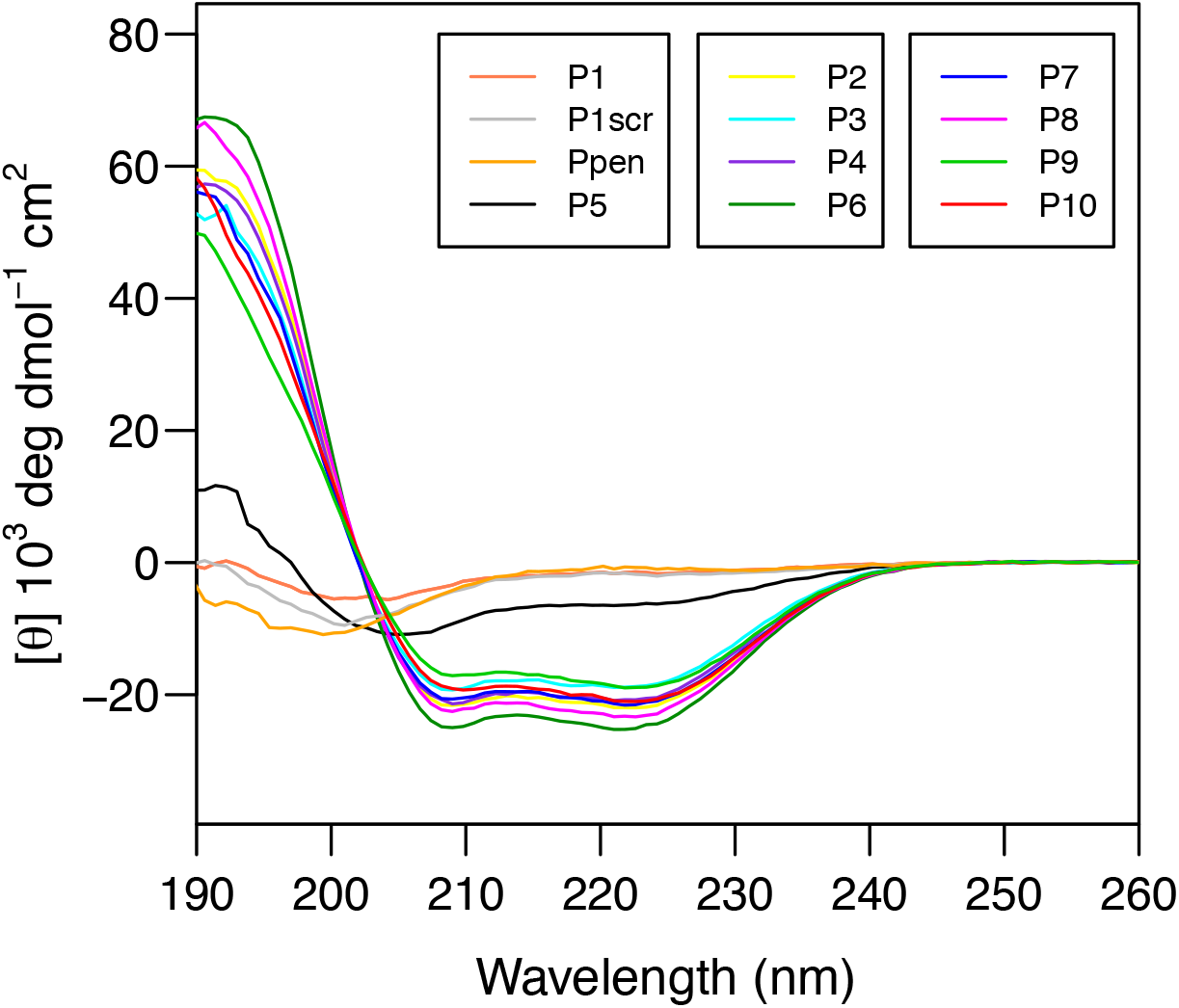
Far-UV CD spectra of synthesized peptides (60 μM in 50 mM sodium phosphate buffer at pH 7.4). CD measurements are reported as molar ellipticity per residue.

### Peptide mimics of hACE2 show high anti-infective efficacy and are devoid of cell toxicity

To determine whether our peptide mimics of hACE2 **H1** helix block SARSCoV-2 cell infection, antiviral assays^25^ were performed on VeroE6 cell line (ATCC CRL-1586) (**Figure 3**) with SARS-CoV-2 clinical isolate obtained from Bronchoalveolar lavage (BAL) of a symptomatic infected patient (#SARS-CoV-2/PSL2020) at Pitié-Salpêtrière hospital, Paris (France) (See M). We first measured inhibition of viral replication in VeroE6 cell cultures exposed to 10 μM of the first set of peptide mimics (**P2** to **P8**, **P1**, **P1scr** and **Ppen** being used as controls), for 48h (Figure **3A**). These preliminary assays helped us to identify two peptide mimics that stand out (**P7** and **P8**) for their ability to block the viral infection, highlighting a potential role of homotyrosine 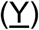. This observation helped us in the peptide mimics structure optimization process. Two new peptides were designed, **P9** and **P10** incorporating the hTyr residue and evaluated with **P8** for their ability to block viral infection on VeroE6 cells through the measurement of infectious virus production and viral genome.^26^ In order to get insight on their ability to block viral infection on human pulmonary cells, Calu3 cell line (ATCC HTB55) was chosen. This pulmonary epithelial cell line acts as respiratory models in preclinical applications^27^ on which SARS-CoV-2 replicated efficiently.^28^

**Figure 3.**
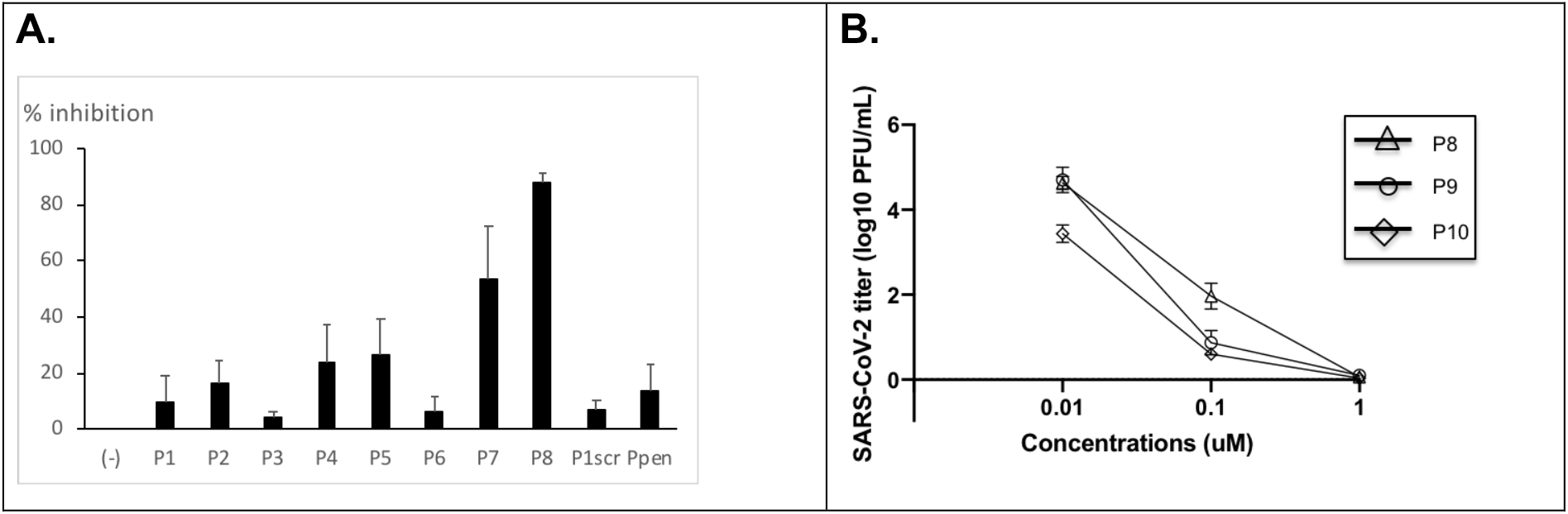
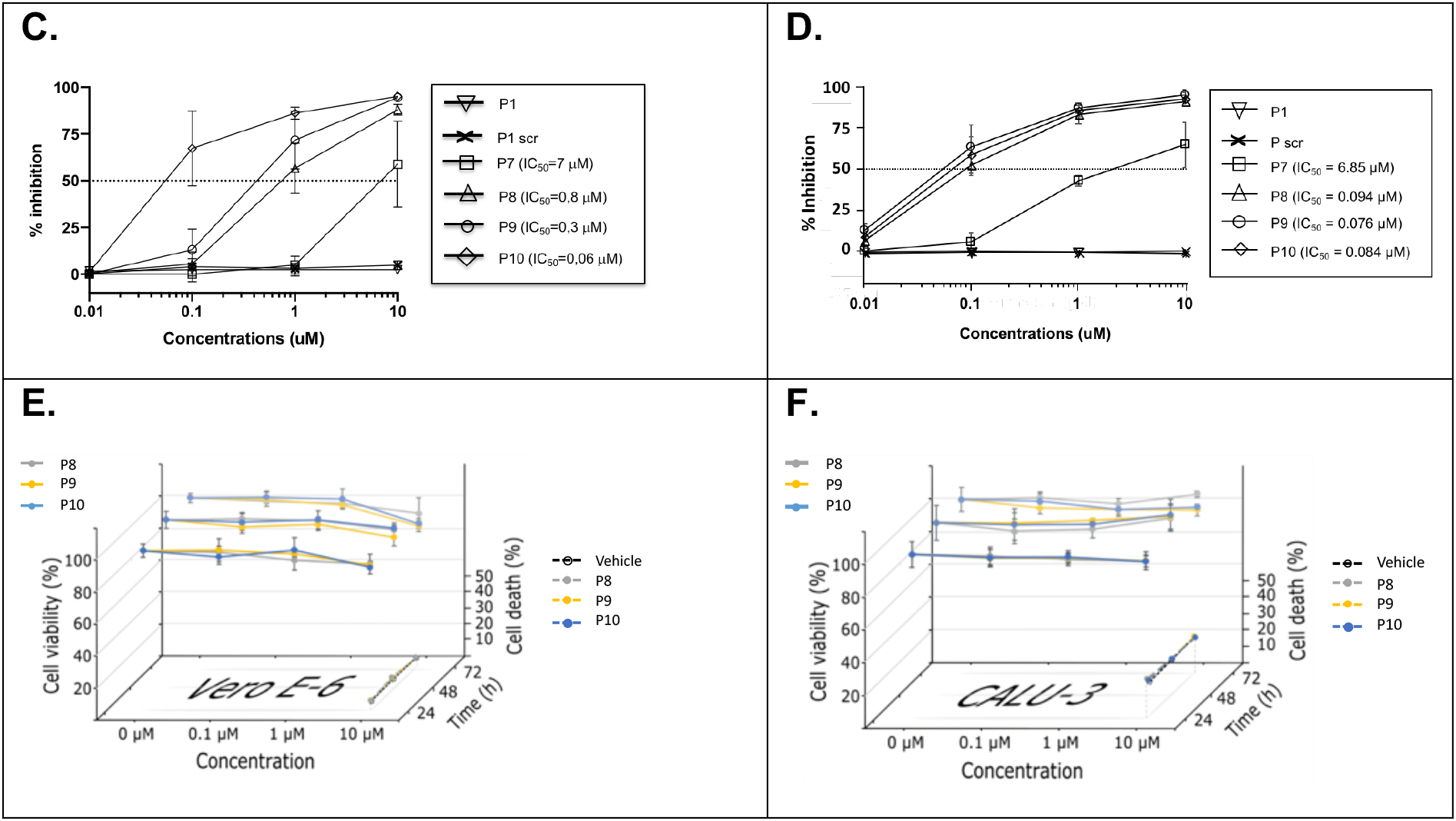
A peptide mimic of hACE2 shows high anti-infective efficacy and is devoid of cell toxicity. **A.** Percent inhibition of SARS-Cov-2 replication. Vero cells were infected with SARS-CoV-2/PSL2020 P2 stock at a multiplicity of infection (MOI) of 0.5 in the presence of 10 peptides (**P1 to P8, P1scr and Ppen as controls)**) at 10 μM for 2 hr, after which the virus was removed and cultures were washed, incubated for 48 h, before supernatant was collected to measure virus replication by Elisa. Data are combined from 3 to 6 independent experiments and expressed as compared to untreated SARS CoV-2-infected Vero cells. The black line is at the median. **B.** SARS-CoV-2 titer reduction. Vero cells were infected with SARS-CoV-2/PSL2020 P2 stock in triplicate at a multiplicity of infection (MOI) of 0.5 in the presence of different concentration (from 0.01 to 10 μM) of peptides **P8**, P9 and **P10** for 2 hr, after which the virus was removed, and cultures were washed in, incubated for 72 h to measure virus production by plaque assay. **C** and **D.** Concentration dose-responses. VeroE6 cells (**3C**) and Calu3 cells (**3D**) were infected with SARS-CoV-2/PSL2020 P2 stock at respectively a multiplicity of infection (MOI) of 0.1 and 0.3 in the presence of different concentration (from 0.01 to 10 μM) of peptides **P1**, **P1 scr**, **P7**, **P8, P9, P10**, for 2 hr, after which the virus was removed, and cultures were washed in, incubated for 48 h, before supernatant was collected to measure virus replication by Elisa. Data are combined from 3 to 6 independent experiments and expressed as percent of inhibition compared to untreated SARS CoV-2-infected Vero cells. Dotted line at 50 % inhibition served to the measure of the IC50. **E** and **F.** Cell viabilities of VeroE6 cells (**3E**) and Calu3 cells (**3F**) were measured by MTT assays after treatment with vehicle 0, 0.1, 1 or 10 μM of P8, P9 or P10 for 24, 48, or 72 h. Cell death was measured by flow cytometry using annexin-V-APC and PI staining in CALU-3 or Vero-E6 cells treated with vehicle or 10 μM P8, P9 or P10 for 24, 48, or 72 h. The plots represent the means (+-SD) of three independent experiments.

We first observed a dose-dependent reduction in virus titer (Figure **3B**) and then calculated average median inhibitory concentration (IC_50_) on VeroE6 and Calu3 cells for **P8**, **P9** and **P10**, with respectively (IC_50_) of 800 nM, 300 nM and 60 nM (Figure **3C**) and 94 nM, 76 nM and 84 nM (Figure **3D**). Importantly, no cytotoxicity was observed in similarly treated uninfected culture cells at 10 μM (concentration more than 150 times higher than IC_50_ for the most potent peptide mimics, Figure **3E** and **3F** and SI). Collectively, these data demonstrate the high antiviral potency of peptide analogs **P9** and **P10**.

### The designed peptides bind to SARSCoV-2 spike RBD with high affinity

Finally, the peptides able to block the cell infection with IC_50_ in the nM-sub μM range (**P8**, **P9** and **P10**) were evaluated for their ability to bind to SARSCoV-2 spike RBD (Figure 4) using Biolayer Interferometry (BLI) with an Octet RED96e (FortéBio).^29^ hACE2 and **P1** were used as respectively positive and negative controls.

**Figure 4.**
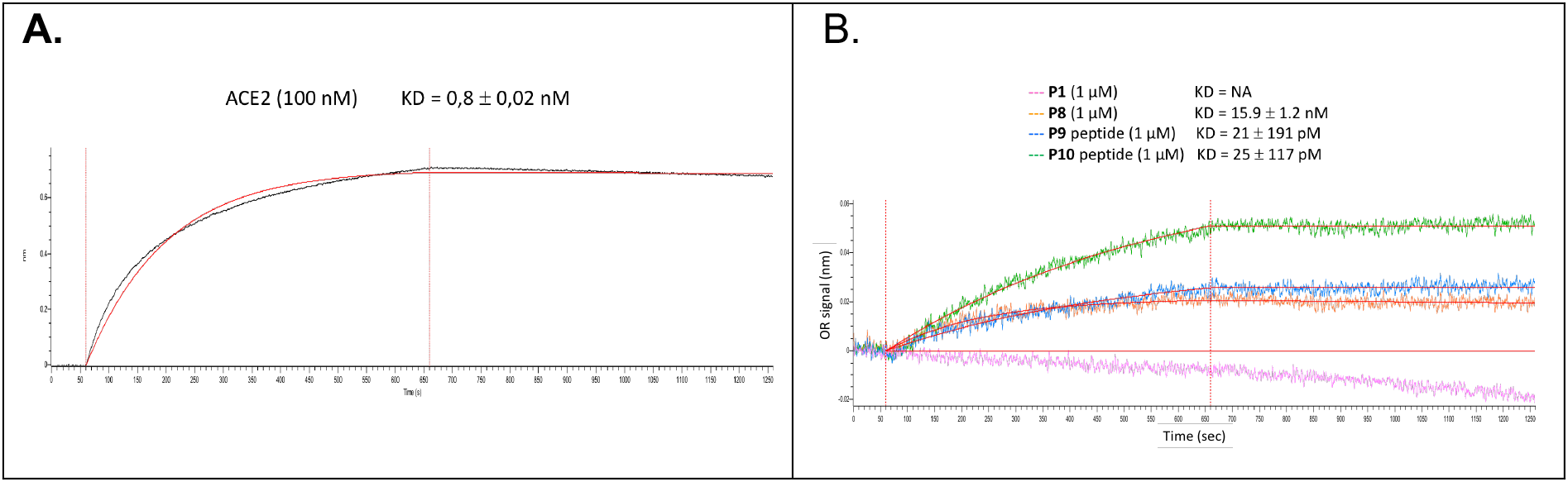
BLI binding experiments. Peptide **P1** and hACE2 were used as a negative and positive controls, respectively. The Fc-tagged 2019-nCoV RBD-SD1 (Sanyou Biopharmaceuticals Co. Ltd) was immobilized to an anti-human capture (AHC) sensortip (FortéBio) using an Octet RED96e (FortéBio). The sensortip was then dipped into 100 nM hACE2 (Sanyou Biopharmaceuticals Co. Ltd, His Tag) (Figure **4A**) or 1 μM of any tested peptide (**Figure 4B**) to measure association before being dipped into a well containing only running buffer to measure dissociation. Data were reference subtracted and fitted to a 1:1 binding model using Octet Data Analysis Software v11.1 (FortéBio) and reported in table 1 as the Kd value.

Even though this technique presents some drawbacks and at least poor sensitivity when the immobilized sample is a protein of high molecular weight and the binding partners are peptides, it remained useful to identify and rank binding peptides. Of note, only association rates could be quantified accurately, dissociation ones being very slow, highlighting strong binding properties of some peptides. Data indicate that the most efficient peptides **P9** and **P10** bind with estimated Kd below 1 nM, whereas **P1** does not bind to spike RBD in the conditions tested here.

## Discussion

The pandemic caused by SARS-CoV-2 is at the origin of an unprecedented health crisis. The medical world has found itself helpless in the face of this virus, having to deal with the absence of specific effective treatment. To date, clinically approved vaccines or specific drugs addressing SARS-CoV-2 targets are lacking.^30^ Among all possible viral targets, the virus spike protein/hACE2 interaction has been validated and the design of compounds able to block this interaction upon binding to spike protein is a promising approach. However, developing a specific drug at a pandemic speed is a hard task and this specific drug cannot be a small molecule. Indeed, beyond the time required for the identification and validation of a lead compound after a library screening, followed by structure-activity relationship studies and clinical development, small molecule drugs are associated with a high attrition rate partly due to their off-target toxicity observed during pharmacological studies.

Peptides appear here as a possible solution for design and development at pandemic speed. Peptides are widely recognized as promising therapeutic agents for the treatment of various diseases such as cancer, and metabolic, infectious, or cardiovascular diseases^31,32^ Across the world and to date, about 70 peptide drugs have reached the market and 150 are currently under clinical development.^28,29^ Special advantages that peptides show over other drugs include their high versatility, target-specificity, lower toxicity, and ability to act on a wide variety of targets^33^ which are directly responsible for greater success rate than small molecules (approval rate of around 20% versus 10%)^34,35^. The syntheses and the development of long therapeutic peptides (over 30 residues) are no more a challenge, as highlighted by the success story of many GLP-1 analogs.^36^ Their possible antigenicity can be evaluated using prediction tools in the design.^37^

Of course, even for peptides, the development of a drug at a pandemic speed requires some considerations. Our aim was to design a peptide with reasonable helical folding propensity in water in order to mimic the **H1** helix of hACE2 in the protein context, considering this helical folding as a prerequisite to compete with hACE2 upon interaction with viral spike protein. The design was realized using only natural amino acids^38^ and avoiding complex tools known and validated to stabilize α-helix.^39,40^ Stabilizing α-helical structure of medium size peptide sequences (up to 15 residues) using only natural amino acids is a hard but achievable task.^41^ Our choice was guided by the desire to build a simple peptide easy to produce quickly on a large scale, without technical constraints requiring sometimes laborious development. We also assumed that the use of mostly natural amino acids can facilitate the essential stages of the development of therapeutic tools in the event of success, particularly around pharmacokinetics, pre-clinical and clinical toxicity aspects. This seems to us to be a fair compromise between designing α-helix peptide with optimized binding affinity and developing an effective tool within short deadlines by integrating the constraints of developability.

Using a combination of validated methods, we improved the helical folding propensity of the native α-helix extracted from the protein context, thanks to leucine and alanine scanning (See tables **1**, **S1** and **S2**), designing and synthesizing a first set of peptides (**P1** to **P8**). These peptides demonstrated to have a high helical content (up to 80% for **P6**). However, increasing this helical content to maximum level led to increasing mean hydrophobicity and hydrophobic moment (see **Table S4**) that proved to be detrimental to solubility and efficacy. The substitution of a leucine residue by the *homo*Tyrosine residue led to peptide analogue **P7** with a slight increase in solubility and a weak efficiency to block SARS-CoV-2 cell infection (IC_50_=7 μM). In this first generation of peptides, the 27 residue **P8** peptide appeared to be highly soluble with high helical folding propensity (70%), and an ability to block SARS-CoV-2 cell infection at 10 μM with an IC_50_ of 800 nM on VeroE6 cells.

In order to improve the potency of our peptides, we designed a new set of mimics combining the properties of peptides **P7** and **P8**, *i.e.* **P9** and **P10**. If the Leu/hTyr substitution led to a slight decrease of experimental helical content, this was at the advantage of the mean hydrophobicity (see **Table S4**) also highlighted by lower HPLC retention times (see **Table S3**). These peptides proved to be highly efficient in blocking SARSCoV-2 cell infection (100% efficacy at 1 μM) on VeroE6 cells together on pulmonary cells with IC_50_ in the nM ranges. This blocking property is related to their ability to bind to SARSCoV-2 spike RBD with affinity estimated in the sub-nanomolar range (**Tables 1** and **S6**). Finally, these peptides proved to be devoid of cell toxicity at 150 times the IC_50_ concentration (**Figures 3E and F** and SI) highlighting their therapeutic potential.

## Conclusions and perspectives

We demonstrated here the feasibility of designing hACE2 peptide mimics with high helical folding propensity in water. The folding propensity promotes interaction with spike RBD and blocks SARS-CoV-2 pulmonary cell infection. Devoid of cell toxicity even at high doses, these mimics might be considered for prophylactic or therapeutic purposes upon adequate formulation. Targeting prophylaxis first might shorten the drug development time scale. Formulated as a sublingual tablet or oral spray, these peptides might be aimed at blocking the infectivity of the virus in a preventive manner. Their biodistribution would be limited to the upper airways (oral cavity …) and it would be degraded in the digestive tracks without any toxic residues.

## Funding and Acknowledgements

This work was supported by private funds (PK), SATT-Lutech, Kaybiotix (LGM PhD grant), French Research Ministry (EO PhD grant). PK is grateful to SATT-Lutech team for its flawless support from the start of this project, to Fabrice Viviani and Akanksha Gangar from Oncodesign for their unwavering support. The authors thank David Boutolleau from the Virology Department, Pitié-Salpêtrière, AP-HP, were was diagnosed the BAL SARS-COV-2 infection.

## Authors Contributions

PK conceived and supervised this project, designed and synthesized the peptides, designed the experiments, interpreted the data and wrote the draft and the discussion of the manuscript. PK and OL wrote the manuscript. PK and OL performed the molecular modeling study. AD contributed to the molecular modeling study. OL performed the CD structural studies. VV performed the cell inhibition assays. PG performed the binding experiments. EO performed the peptides LC-MS analyses. LGM performed the cell toxicity experiments. AG and CEL recruited the patient and provided BAL sample.

## Abbreviations Used

SARS(-CoV-2), severe acute respiratory syndrome (-coronavirus 2); COVID-19, coronavirus disease 2019; hACE2, human angiotensin converting enzyme 2. AD, antigenic determinant; LCMS, liquid chromatography mass spectroscopy. CD, Circular Dichroism.

## Methods

### 1. General chemistry

#### 1.1 Peptides syntheses

Peptides were produced manually synthesized from Fmoc-protected amino acids utilizing standard solid phase peptide synthesis (SPPS) methods. Solid-phase peptide syntheses were performed in polypropylene Torviq syringes (10 or 20 mL) fitted with a polyethylene porous disk at the bottom and closed with an appropriate piston. Solvent and soluble reagents were removed through back and forth movements. The appropriate protected amino acids were sequentially coupled using PyOxim/Oxyma as coupling reagents. The peptides were cleaved from the rink amide resin with classical cleavage cocktail TFA/TIS/H_2_O (95:2.5:2.5). The crude products were purified using preparative scale HPLC. The final products were characterized by analytical LCMS. All tested compounds were TFA salts and were at least 95% pure. The relevant peptides after CD spectra analyses were selected and produced by Genecust France on 20mg scale.

#### 1.2. Purification

Preparative scale purification of peptides was performed by reverse phase HPLC on a Waters system consisting of a quaternary gradient module (Water 2535) and a dual wavelength UV/visible absorbance detector (Waters 2489), piloted by Empower Pro 3 software using the following columns: preparative Macherey-Nagel column (Nucleodur HTec, C18, 250 mm × 16 mm i.d., 5 μm, 110 Å) and preparative Higgins analytical column (Proto 200, C18, 150 mm × 20 mm i.d., 5 μm, 200 Å) at a flow rate of 14 mL/min and 20 mL/min, respectively. Small-scale crudes (<30 mg) were purified using semipreparative Ace column (Ace 5, C18, 250 mm × 10 mm i.d., 5 μm, 300 Å) at a flow rate of 5 mL/min. Purification gradients were chosen to get a ramp of approximately 1% solution B per minute in the interest area, and UV detection was done at 220 and 280 nm. Peptide fractions from purification were analyzed by LC−MS (method A or B depending of retention time) or by analytical HPLC on a Dionex system consisting of an automated LC system (Ultimate 3000) equipped with an autosampler, a pump block composed of two ternary gradient pumps, and a dual wavelength detector, piloted by Chromeleon software. All LC−MS or HPLC analyses were performed on C18 columns. The pure fractions were gathered according to their purity and then freeze-dried using an Alpha 2/4 freeze-dryer from Bioblock Scientific to get the expected peptide as a white powder. Final peptide purity (>95%) of the corresponding pooled fractions was checked by LC−MS using method A.

#### 1.3. Analytics

Two methods were conducted for L1C4–MS analysis.

##### Method A

Analytical HPLC was conducted on a X-Select CSH C18 XP column (30 mm × 4.6 mm i.d., 2.5 μm), eluting with 0.1% formic acid in water (solvent A) and 0.1% formic acid in acetonitrile (solvent B), using the following elution gradient: 0−3.2 min, 0−50% B; 3.2−4 min, 100% B. Flow rate was 1.8 mL/min at 40 °C. The mass spectra (MS) were recorded on a Waters ZQ mass spectrometer using electrospray positive ionization [ES+ to give (MH)+ molecular ions] or electrospray negative ionization [ES− to give (MH)− molecular ions] modes. The cone voltage was 20 V.

##### Method B

Analytical HPLC was conducted on a X-Select CSH C18 XP column (30 mm × 4.6 mm i.d., 2.5 μm), eluting with 0.1% formic acid in water (solvent A) and 0.1% formic acid in acetonitrile (solvent B), using the following elution gradient: 0−3.2 min, 5−100% B; 3.2−4 min, 100% B. Flow rate was 1.8 mL/min at 40 °C. The mass spectra (MS) were recorded on a Waters ZQ mass spectrometer using electrospray positive ionization [ES+ to give (MH)+ molecular ions] or electrospray negative ionization [ES− to give (MH)− molecular ions] modes. The cone voltage was 20 V.

### 2. CD Spectroscopy

CD spectra were recorded on a Jasco J-815 CD spectropolarimeter equipped with a Peltier temperature controller. Data were obtained at 25°C over a wavelength range between 185 and 270 nm, using a wavelength interval of 0.2 nm and a scan rate of 20 nm/min. Peptide samples were prepared at a concentration of 60 μM in 50 mM sodium phosphate buffer, pH 7.4, in a quartz cell of 1 mm path length. CD experiments were processed and plotted with R program. CD spectra were analyzed using DICHROWEB web server and CDSSTR deconvolution algorithm.^24^

### 3. Anti-infectivity study on Vero-E6^25^ and Calu3 cells

#### 3.1. Cells and virus preparation

VeroE6 (ATCC CRL-1586) and Calu3 (ATCC HTB55) cells were cultured in Dulbecco’s Modified Eagle Medium (DMEM) supplemented with non-essential amino acids (NEAA), penicillin/streptomycin (P/S), and 10% (v/v) fetal bovine serum (FBS). SARS-CoV-2 clinical isolate was obtained from Bronchoalveolar lavage (BAL) of a symptomatic infected patient (#SARS-CoV-2/PSL2020) at Pitié-Salpêtrière hospital, Paris (France). BAL (0.5 mL) was mixed with an equal volume of DMEM without FBS, supplemented with 25mM Hepes, double concentration of P/S and miconazole (Sigma), and added to 80% confluent Vero-E6 cells monolayer seeded into a 25 cm^2^ tissue culture flask. After 1 h adsorption at 37°C, 3 mL of infectious media (DMEM supplemented with 2% FBS, P/S and miconazole) were added. Twenty-four hours post-infection another 2 mL of infectious media were added. Five days post-infection, supernatants were collected, aliquoted and stored at −80°C (P1). For secondary virus stock, Vero-E6 cells seeded into 25 cm^2^ tissue culture flasks were infected with 0.5 mL of P1 stored aliquot, and cell-culture supernatant were collected 48 h post-infection, and stored at −80°C (P2). Infectious viral particles were measured by a standard plaque assay previously described with fixation of cells 72 hr post infection, as described (Mendoza et al, 2020). Accordingly, the viral titer of SARS-CoV-2/PSL2020 P2 stock is about 5.3 10^5^ PFU/mL.

#### 3.2. Experiments of peptide-neutralization

VeroE6 or Calu3 (1 × 10^5^ cells/mL) were seeded into 24 wells plates in infectious media and treated with different concentrations of the peptides (from 0.1 to 10 μM). After 30 min at room temperature, cells were infected with 0.1 moi (VeroE6) or 0.3 moi (Calu3) of SRAS-Cov2 (SARS-CoV-2/PSL2020 P2 stock) in infectious media. All conditions were tested in triplicate.

Cell supernatants were collected at 48 h post-infection for Elisa assay using SARS-CoV-2 (2019-nCoV) Nucleoprotein / NP ELISA Kit from Sino biological, according the manufacturer’s instruction, and standard plaque assay.

### 4. Toxicity study on Vero-E6 cells and Calu3

Cell viability was measured by MTT assays after treatment with 0, 0.1, 1.0 or 10 μM of the indicated peptide for 24, 48, or 72 h.

Methodology: 96-well plates were used to seed 10^4^ CALU-3 or Vero-E6 cells per well, which were let to adhere. Vero-E6 cells reached ~50% confluency the day after, while CALU-3 did it after 48 h. When this was reached, the medium in the wells was replaced and cells were either left alone (0 μM) or treated with 0.1 μM, 1 μM or 10 μM of the corresponding peptide. MTT (2 mM) was added to each well after 24 h, 48 h or 72 h of treatment and incubated 4 h in a controlled, humidified atmosphere with 5% CO_2_ at 37 ºC. Supernatant in each well was discarded from the wells using a vacum pump and formazan salts were disolved in 100 μL DMSO to read plate absorbance at 570 nM. Absorbance in each well was normalized with untreated controls. The plots represent the means (± SD) of an experiment performed in triplicates

### 5. 5-Biolayer Interferometry experiments

Fc-tagged 2019-nCoV RBD-SD1 (Sanyou Biopharmaceuticals Co. Ltd) was immobilized to an anti-human capture (AHC) sensortip (FortéBio) using an Octet RED96e (FortéBio). The sensortip was then dipped into 100 nM hACE2 (Sanyou Biopharmaceuticals Co. Ltd, His Tag) or 1 μM of any tested peptide to measure association before being dipped into a well containing only running buffer composed of DPBS (Potassium Chloride 2.6mM, Potassium Phosphate monobasic 1.5mM, Sodium Chloride 138mM, Sodium Phosphate dibasic 8mM), 0.05% Tween 20 and 0.5% bovine serum albumin to measure dissociation.

Data were reference subtracted and fit to a 1:1 binding model using Octet Data Analysis Software v11.1 (FortéBio) and reported on figure **4**.

